# Quantifying Evoked Responses through Information-Theoretical Measures

**DOI:** 10.1101/2022.11.11.516096

**Authors:** Julian Fuhrer, Kyrre Glette, Anaïs Llorens, Tor Endestad, Anne-Kristin Solbakk, Alejandro Blenkmann

**Affiliations:** RITMO Centre for Interdisciplinary Studies in Rhythm, Time and Motion, University of Oslo, Oslo, Norway; Department of Informatics, University of Oslo, Oslo, Norway; Department of Psychology, University of Oslo, Oslo, Norway; Helen Wills Neuroscience Institute and Department of Psychology, University of California, Berkeley, USA; Department of Neurosurgery, Oslo University Hospital, Rikshospitalet, Oslo, Norway; Department of Neuropsychology, Helgeland Hospital, Mosjøen, Norway

**Keywords:** EEG, SEEG, ECOG, Information Content, Algorithmic Complexity, t-test, Frequency Tagging, Mutual Information

## Abstract

Information theory is a viable candidate to advance our understanding of how the brain processes information generated in the internal or external environment. With its universal applicability, information theory enables the analysis of complex data sets, is free of requirements about the data structure, and can help infer the underlying brain mechanisms. Information-theoretical metrics such as Entropy or Mutual Information have been highly beneficial for analyzing neurophysiological recordings. However, a direct comparison of the performance of these methods with well-established metrics, such as the t-test, is rare. Here, such a comparison is carried out by evaluating the novel method of Encoded Information with Mutual Information, Gaussian Copula Mutual Information, Neural Frequency Tagging, and t-test. We do so by applying each method to event-related potentials and event-related activity in different frequency bands originating from intracranial electroencephalography recordings of humans and marmoset monkeys. Encoded Information is a novel procedure that assesses the similarity of brain responses across experimental conditions by compressing the respective signals. Such an information-based encoding is attractive whenever one is interested in detecting where in the brain condition effects are present.

## Introduction

Efficient processing of *information* is a core capacity of the brain. It enables us to perceive rapidly, comprehend, identify changes, and engage with our environment – sometimes even without conscious effort. Accordingly, in attempting to understand the underlying brain mechanisms responsible for these capacities, it is useful to employ approaches that originate from *information* theory. This mathematical theory provides multivariate analysis tools, is not bound to a single type of data, is model-independent (i.e., does not require assumptions about the data itself), and can capture nonlinear interactions (1–4). Specifically, by measuring the degree of redundancy, the branch of *algorithmic* information theory (AIT) estimates the absolute information contained in individual brain responses. The higher the information content, the more complex its structure. Accordingly, the less compressible or more random the response (2). Assessing the absolute information content of responses recorded from different contact sites enables inferring the complexity of the activity recorded at the respective sensors and potentially identification of underlying dynamics.

Hence, principles from information theory can be utilized for the analysis of neurophysiological recordings originating from scalp electroencephalography (EEG), intracranial EEG (iEEG), magnetoencephalography (MEG), or functional magnetic resonance imaging (fMRI). While information theory-based metrics have been employed to advance our understanding of brain processes, direct comparisons with well-established metrics or demonstrations of their easy applicability are highly needed. More specifically, AIT has been applied to analyses of task-related cognitive operations or to discriminate between states of consciousness measured with EEG, iEEG, MEG, or fMRI recordings (5–9), however, its use in neuroscience has been limited despite its clear potential.

Here, we carry out such a comparison and demonstrate the AIT-based measure of *encoded information* (EI; 10) as an advantageous tool to directly quantify the level of similarity between responses of neurophysiological data across experimental conditions. We do so by analyzing iEEG recordings stemming from 34 humans (11, 12) and three marmoset monkeys (13, 14) which were exposed to passive and active paradigms as well as auditory or visual stimuli. We compare the performance of encoded information with that of a conventional t-test, Mutual Information (MI), Gaussian Copula Mutual Information (GCMI; 3), or Neural Frequency Tagging (NFT; 15, 16) considering the signal band-pass power time series of theta (5 to 7 Hz), alpha (8 to 12 Hz), beta (12 to 24 Hz), high-frequency (HFA; 75 to 145 Hz), and event-related potential (ERP).

## Material & Methods

### Test Paradigms and Neurophysiological Recordings

To evaluate the methods, we examined their sensitivity to discriminate experimental conditions from three different neurophysiological data sets. Two of these used auditory stimuli and one used visually presented stimuli. We focused our analysis on the cortical representation of theta, alpha, beta, HFA, and ERP band-pass power time series.

### Extraction of Band-pass Power Time Series

Data were low-pass filtered at 30 Hz using a sixth-order Butterworth filter to obtain the ERPs. Theta, alpha, and beta frequency bands were extracted from the demeaned signals using wavelet time-frequency transformation (Morlet wavelets) based on convolution in the time domain (17). Wavelets of 3, 3, or 5 oscillations were used to extract respective frequencies bands (theta (5 to 7 Hz), alpha (8 to 12 Hz), and beta (12 to 24 Hz)) in steps of 1 Hz. All trials were then baseline corrected by subtracting the mean amplitude of the baseline period of each trial and frequency band from the entire trial (see respective sections for the different data sets for the used baseline intervals).

To extract the HFA, the pre-processed data were filtered into eight bands of 10 Hz ranging from 75 to 145 Hz by use of bandpass filters. Next, the instantaneous amplitude signal of each filtered signal was computed by applying a Hilbert transform to the filtered time series leading to the analytic signal (18), constituting a complex-valued time series. The analytic amplitude time series or signal envelope corresponding to specific frequency bands was then obtained using Pythagoras’ Theorem. To obtain one time series across all eight frequency bands, their mean amplitude value was calculated. As the last step, the respective time series were normalized by dividing them by a mean baseline period computed from all trial recordings. This resulted in a normalized measure relative to the baseline activity and termed HFA.

To eliminate any residual artifacts not rejected by visual inspection, responses with an amplitude larger than five standard deviations from the mean for more than 25 consecutive ms, or with a power spectral density above five standard deviations from the mean for more than six consecutive Hz were excluded.

### Optimum-1 Paradigm

We analyzed experimental iEEG data obtained from intracranial electrodes implanted in (self-reported) normal-hearing adults with drug-resistant epilepsy. Analyses of this data have been previously presented in (5, 10, 11). Participants (n=22, mean age 31 years, range 19 to 50 years, 6 female) performed a passive auditory oddball paradigm where a standard tone alternated with random deviant tones. The tones had a duration of 75 ms and were presented every 500 ms in blocks of 5 min consisting of 300 standards and 300 deviants. At the beginning of each block, 15 standards were played. To capture automatic, stimulus-driven processes, participants were asked not to pay attention to the sounds while reading a book or magazine. They completed 3 to 10 blocks, providing at least 1800 trials (for details, see 10, 11). From the 22 participants, a total of 1078 channels (mean: 48, range: 12 to 104) were available after data cleaning. Data were then segmented into 2000 ms epochs (750 ms before and 1250 ms after tone onset) and demeaned. The different band-pass power time series were then extracted. For each time series, differences between standard and deviant tone responses were then evaluated in the 400 ms time window following the sound onset across channels and subjects. The baseline window was from −100 to 0 ms relative to tone onset. Additionally, for this data set the neural oscillation synchronization to the tone onset frequency (2 Hz) and the frequency for standard and deviants tones (1 Hz) were examined by considering the pre-processed data (needed for the NFT approach).

### Roving Oddball Paradigm

Further, we considered experimental iEEG data collected during a passive auditory Roving Oddball paradigm on three awake adult male common marmosets (*callithrix jacchus*). This experimental data has been previously studied in Komatsu et al. (13), Canales-Johnson et al. (14). The paradigm consisted of trains of three, five, or 11 repeated single tones of 20 different frequencies (250 to 6727 Hz with intervals of 1/4 octaves). All tones were pure sinusoidal tones, lasted 64 ms (7 ms rise/fall) and there was stimulus onset asynchrony 503 ms between them. For each sound train, all tones were identical but varied across tone trains. Consequently, the mismatch occurred between the transition from one train to another in the form of a frequency change. Accordingly, the last tone of a train is defined as a standard tone, while the first tone of a new train is considered a deviant tone. Standard to deviant tone transitions then occurred 240 times during a recording session.

The number of implanted electrodes varied from monkey to monkey. For monkey “Fr”, 32 channels were implanted in the left hemisphere epidural space, for “Go”, 64 channels were implanted in the right hemisphere, and for monkey “Kr”, 64 electrodes were implanted in the right hemisphere (Fig. 5c; see 13, 14, for more detailed information). Recorded data were re-referenced through an average reference montage and epoched into −950 to 2000 ms segments relative to the standard or deviant tone onset. The different band-pass power time series were then extracted and baseline corrected (by use of the −100 to 0 ms time interval relative to tone onsets). For the analysis, all recordings were shortened to the −100 to 350 ms interval relative to sound onsets.

### Verbal Working Memory Paradigm

As a third data set, we investigated iEEG data stemming from the insular cortex during a verbal working memory task (vWM; see 12). Participants (n=12, mean age 31.2 ±11.1 yr, 4 female) performed a recent-probes task, where in each trial a list of five letters was displayed (stimulus duration 500 ms, inter-onset interval 1000 ms) on the computer screen. The letter presentation was followed by 4 s maintenance and a 2 s probe period. During the latter, a probe letter was displayed where the participants had to answer whether the presented probe letter was in the current list (p=0.5). In total, 144 trials were presented to each participant in a pseudo-random order within three blocks (each 10 min).

From the twelve participants, a total of 90 bipolar channels (mean: 7.5, range: ±5.9) were available after data cleaning. Data were then segmented into 16 s epochs (−12 s before and 4 s after probe period) and demeaned. The different band-pass power time series were then extracted, and for each band-pass power time series, differences between maintenance (−2 to 0 s) and probe period (0 to 2 s) were evaluated across channels and participants. The window for the baseline correction was from −9.5 to −8.5 s, i.e., the second preceding the presentation of the letter list.

### Encoded Information

We estimated the EI of the mean responses of the different experimental conditions as described in (10). In short, by employing algorithmic or Kolmogorov Complexity (K-complexity), we estimated the EI between conditions through the measure of Normalized Compression Distance (NCD) (2, 19). For a pair of signals (*x, y*), it is defined as

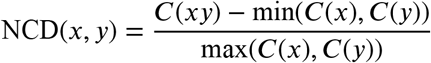

with *C*(*xy*) denoting the compressed size of the concatenation of *x* and *y*, and *C*(*x*) and *C*(*y*) their respective size after compression (2, 19). Further, the NCD is non-negative, that is, it is 0 ≤ NCD(*x, y*) ≤ 1 +ϵ, where the ϵ accounts for the imperfection of the employed compression technique. Small NCD values suggest similar signals and high values indicate rather different signals.

To obtain a compressed version of a signal, it was first simplified by grouping its values into 128 regular intervals (bins) while keeping the temporal sampling rate unchanged. The bins covered equal distances and in a range between the global extrema of all the signals considered. The discrete signal was then compressed by passing an integer representation of the signal to the compressor. This representation constituted the mapping between each respective signal value and the number of the closest bin (6, 7). For instance, the signal value X(*t*) at time point *t* is closest to the bin Q ∈ ℕ with 1 ≤ *Q* ≤ 128. The value for the time point t which is then used for the integer representation is Q. Compression of this integer representation subsequently proceeded through a compression routine based on Python’s standard library with gzip. The statistical hypothesis testing was then performed through a permutation-based approach as described below.

### t-test

The t-test is one of the most common methods to compare amplitude or power time series mean value differences across experimental conditions. For each sample and channel, a two-sided t-test was performed, where the resulting t-value represented the activity difference between the two conditions (e.g., standard or deviant). To correct for multiple comparisons across samples and channels, a False Discovery Rate (FDR) adjustment was applied with an FDR of 0.05.

### Neural Frequency Tagging

For the Optimum-1 data, we used NFT to identify the brain’s capability to automatically segment the continuous auditory stream (15, 16). In the respective auditory sequence, the two main segments are represented by the frequency of sound onsets (*∼*2 Hz) or by the frequency of transitions between standards to standards or deviants to deviants (i.e., half the frequency or *∼*1 Hz). If the neural activity of a recording site showed such “tagging” of frequency-specific properties within the stimuli, we defined it as “responsive”. If it tagged half the frequency, we identified it as sensitive to the pattern of a standard-deviant alternation.

To assess this tagging, we computed the power-spectral density (PSD) of the epoched raw data using Welch’s method (Fig. 2; 20). Subsequently, we estimated the signal-to-noise ratio (SNR) of the PSD (21). Here, SNR defined the ratio of power in a given frequency (signal) to the average power in the surrounding frequencies (noise). By doing so, we normalized the spectrum and accounted for the 1/f power decay (20). We then identified significant peaks through a lower threshold consisting of two times the standard deviation. The latter was estimated by computing the median absolute deviation, which was obtained by taking the median SNR multiplied by the constant distribution-dependent scale factor (Fig. 2). In the case of normally distributed observations, it reflects the 50 % of the standard normal cumulative distribution function, leading to a scale factor of 1.4826 (22, 23). Power peaks of harmonics higher than the tone representation rate of 2 Hz are method-related peaks and are thus not regarded.

### Mutual Information

Besides the t-test and NFT, EI was compared to the measure of MI. While it is also an information-theoretic quantity, in contrast to EI it draws on the concept of Shannon entropy (i.e., classic information theory). For a discrete random variable x with N outcomes, the entropy can be defined as

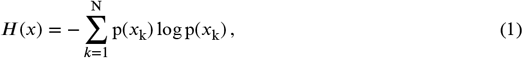

with p(*x*_k_) being the occurrence probability for each element *x*_k_, …, *x*_N_ of *x*. Given this definition, the MI between two discrete random variables (*x, y*) with N or M outcomes can be defined as

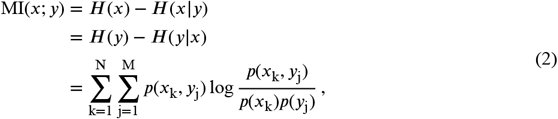

with the joint probability *p*(*x*_k_, *y*_j_) and the marginal probabilities *p*(*x*_k_) and *p*(*y*_j_).

This estimation of MI requires estimating the probability distribution of Eq. 2 by binning the signal into discrete steps. This is followed by a maximum likelihood estimation, yielding the probability distribution estimates. In our analysis, the respective signals were binned into four bins (1, 3).

Besides this binning approach, a novel estimation technique of MI after Ince et al. (3) was employed. With this approach, MI is estimated via Gaussian Copula. In short, each univariate marginal distribution is transformed into a standard normal. Subsequently, a Gaussian parametric MI estimate is applied. That yields a lower bound estimate of MI, named GCMI. Statistical significance testing was then estimated through surrogate data testing.

### Surrogate Testing

The statistical significance of the information-based measures (EI, MI, and GCMI) was assessed through surrogate data testing. Accordingly, p-values were obtained by evaluating the observed information-based quantity in terms of a null distribution (Fig. 1). For EI and binned MI, null distributions were created by repeatedly shuffling the trials (i.e., single evoked responses) between conditions (e.g. standard and deviant) and then re-computing the information-based measure. For the GCMI, this proceeded for each sample of the time series signal. If not noted otherwise, we chose 20 000 iterations to build the null distributions, which is sufficiently above the recommendation of 100 shuffles (24). Single-sided p-values lower or equal to 0.05 were considered statistically significant. To correct for multiple comparisons across channels, FDR adjustment was applied with an FDR of 0.05.

**Figure 1:**
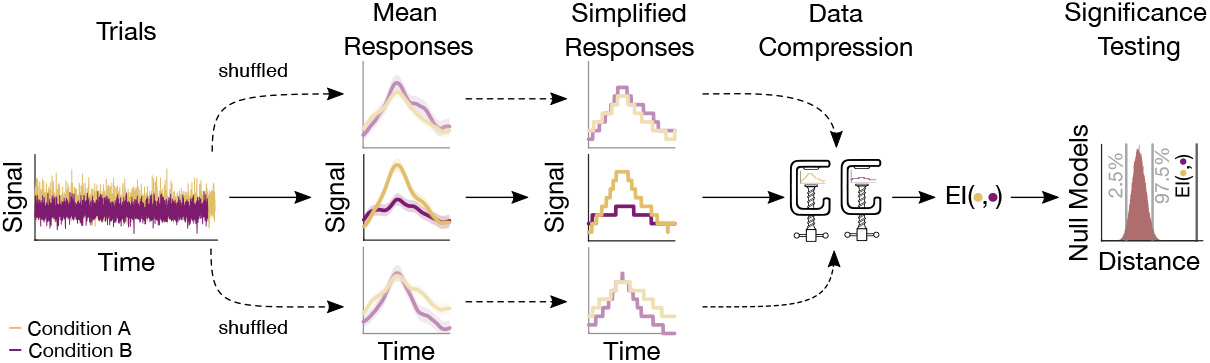
Sketch of the procedure for an example electrode. Based on the trials, a mean response for each condition is computed. Trials are then shuffled, resulting in surrogate mean responses. Subsequently, these signals undergo a simplification procedure, followed by their compression. The output of the compression routine is the EI, quantifying the similarity between responses. The resulting values are then evaluated, leading to a null model distribution. This distribution serves to assess the significance of the actual EI value.

**Figure 2:**
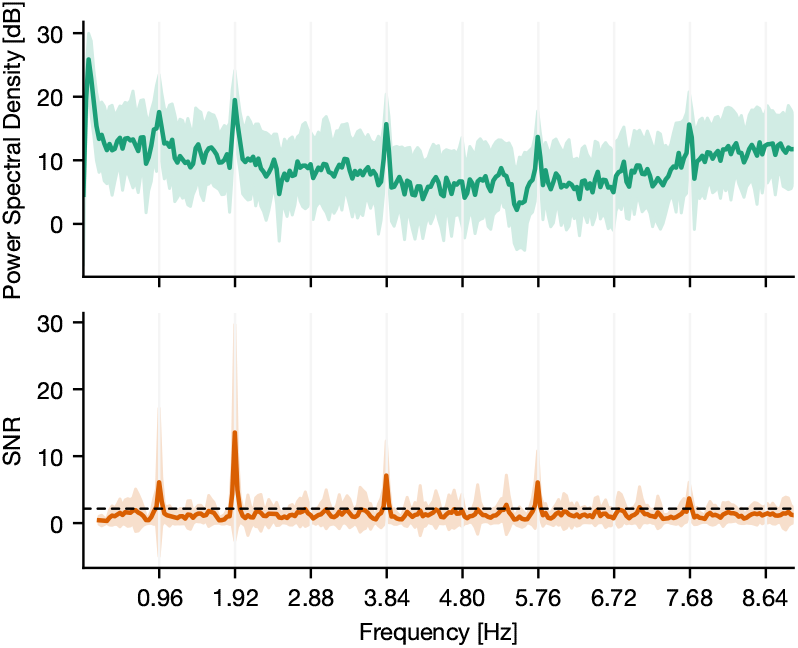
PSD and SNR of a responsive channel located in the superior temporal sulcus of a human. The activity of the channel shows synchrony to the main frequency of tone onsets, which is around 2 Hz. The displayed deviation is due to a constant lag of tone presentation during recording (the resulting theoretical main sequence is at 1.919 Hz). Further, it shows synchrony to half the main frequency indicating that the underlying region of this channel tags the presentation rate of solely standard or deviant tones. Thus, it discriminates between standard and deviant conditions. The dashed line indicates the statistical threshold.

### Significance Ratio

To compare the different methods we determined the number of channels with statistically different activity between conditions relative to the total amount of channels. This ratio was computed either for each subject, where the total significance ratio was the mean ratio across all subjects accompanied by a bootstrapped 95 %-confidence interval. Or by taking the total ratio by collapsing across all channels (regardless of subjects). The latter was used when considering individual brain regions, where the number of channels was limited, leading to distorted ratios with large confidence intervals. For the Roving Oddball data, the ratio of each monkey was considered individually.

### Performance for Limited Amount of Data

We further considered scenarios where only a reduced number of trials were available. Channels were selected from the Optimum-1 paradigm with differing sensitivity to deviating tones. The channels were located in the respective 25, 50, 75, and 97.5-percentiles of the t-value distribution emerging from the t-test-based analysis of HFA (Fig. 7a). The number of trials varied from 1 to 100 % of all available trials (759.50 ± 360.85 trials for deviant responses and 715.14 ± 388.12 trials for standards across all channels). For each percentage and condition, trials were randomly chosen and a null distribution with 1000 surrogates was created. This step was repeated 50 times for each trial increment to obtain an average p-value as a function of the number of trials (Fig. 7b). The surrogate number was chosen in line with the recommendations of Lancaster et al. (24) and the repetition number was based on an empirical approach, keeping it as low as possible to account for the considerable computational resources.

### False Positive Estimates

Evaluating the methods’ performances, we employed neurophysiological data sets and were thus not able to compute the false positive rate of each method given the ground truth. Therefore, we estimated the methods’ false positive rates through a simulation-based approach. The methods were employed to discriminate between two samples drawn from the standard normal distribution (25). Each sample consisted of a time-course signal of 100 observations, was repeatedly drawn (500 times), and for each draw the number of surrogates for EI, MI, or GCMI was 1000. Given that both samples originated from the same distribution, each statistically significant output constituted a false positive. The false positives were then accumulated across simulations and put relative to the respective number of simulations to estimate the false positive rate (Fig. 7c).

## Results

Based on three neurophysiological data sets, we compared the methods’ performances of EI, a conventional t-test, MI, and GCMI on discriminating evoked responses to that across five different cortical band-pass power time series. Additionally, we considered the approach of NFT. Overall, EI and t-test showed the greatest significance ratios (Fig. 3a, 5a, and 6a).

**Figure 3:**
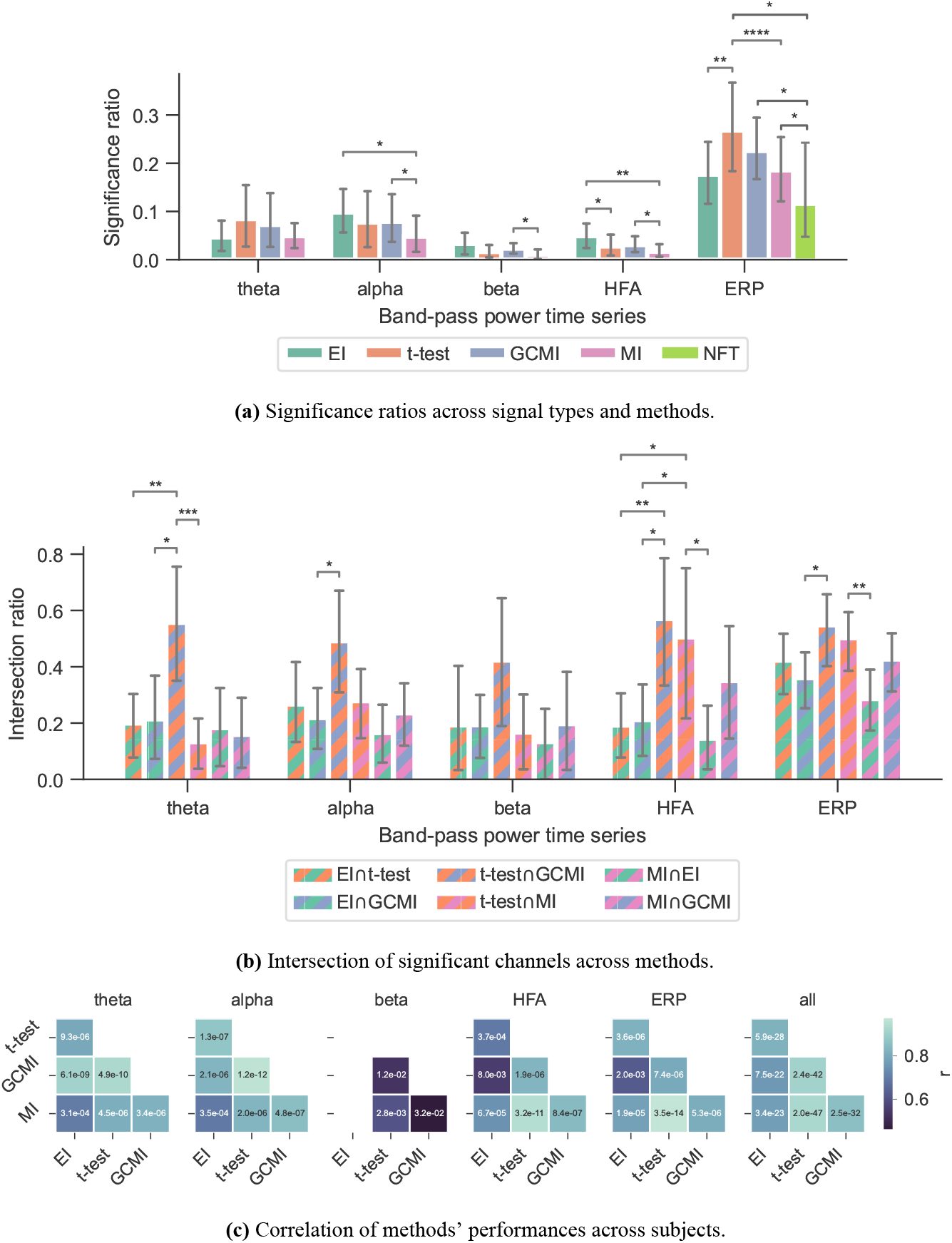
Performance of the different methods for the Optimum-1 paradigm. Error bars indicate 95%-CIs across subjects. Importantly, each subject has a unique electrode distribution such that the range of significant channels can greatly vary. **3a**: Significance ratio for each electrophysiological representation. Statistical significance is indicated with * p≤5*e*−2, ** p≤1*e*−2, *** p≤1*e*−3 and **** p≤1*e*−4. **3b**: Intersection of the significant channels across methods. Each number is shown relative to the total number of significant channels. **3c**: Correlation matrices comparing the subject-specific significant ratios. The respective p-value is annotated in each square.

**Figure 4:**
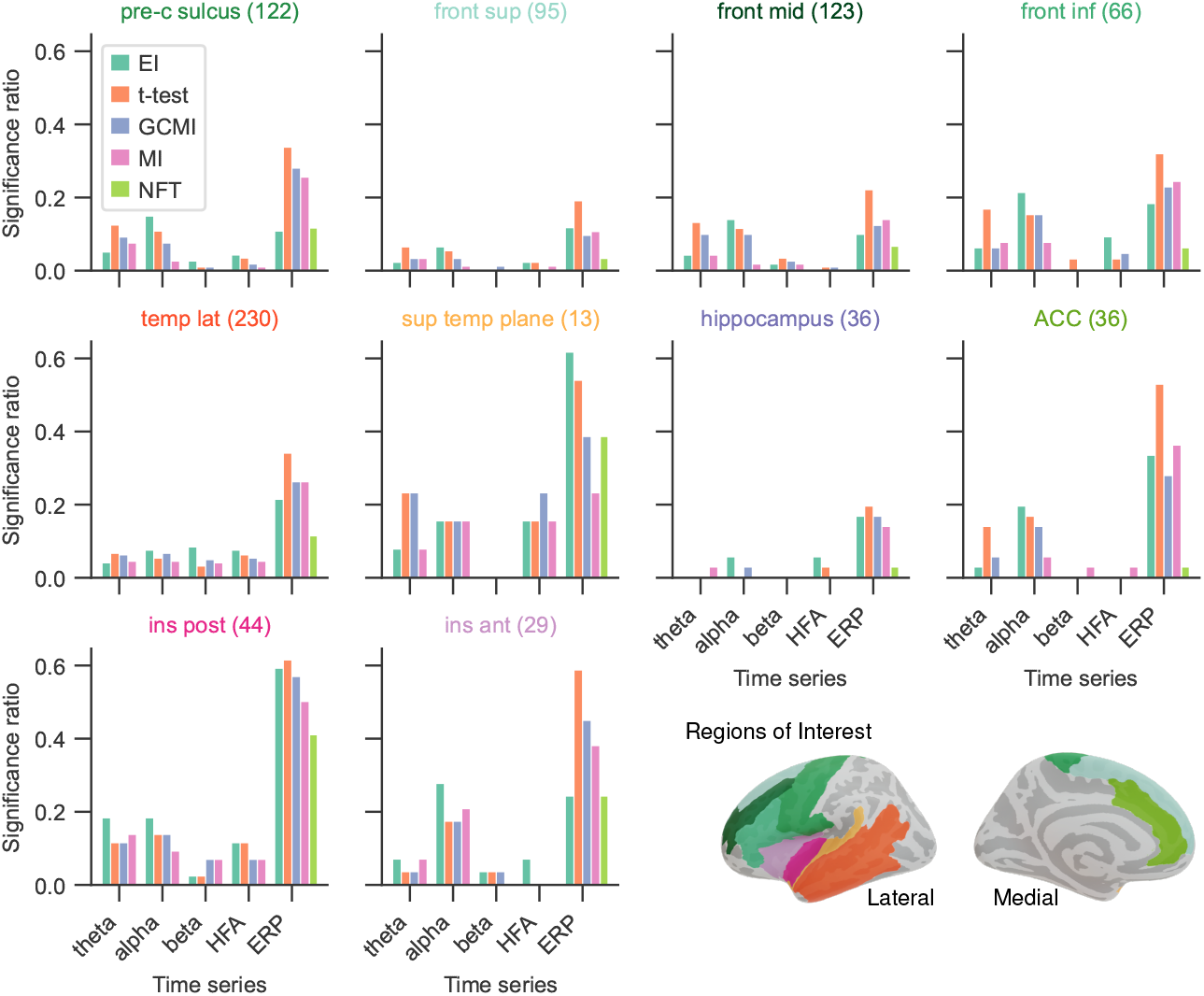
Significance ratio across brain regions for the different methods. The number of channels per ROI is indicated in brackets (For more detailed information on the ROIs, see supplementary of 5).

**Figure 5:**
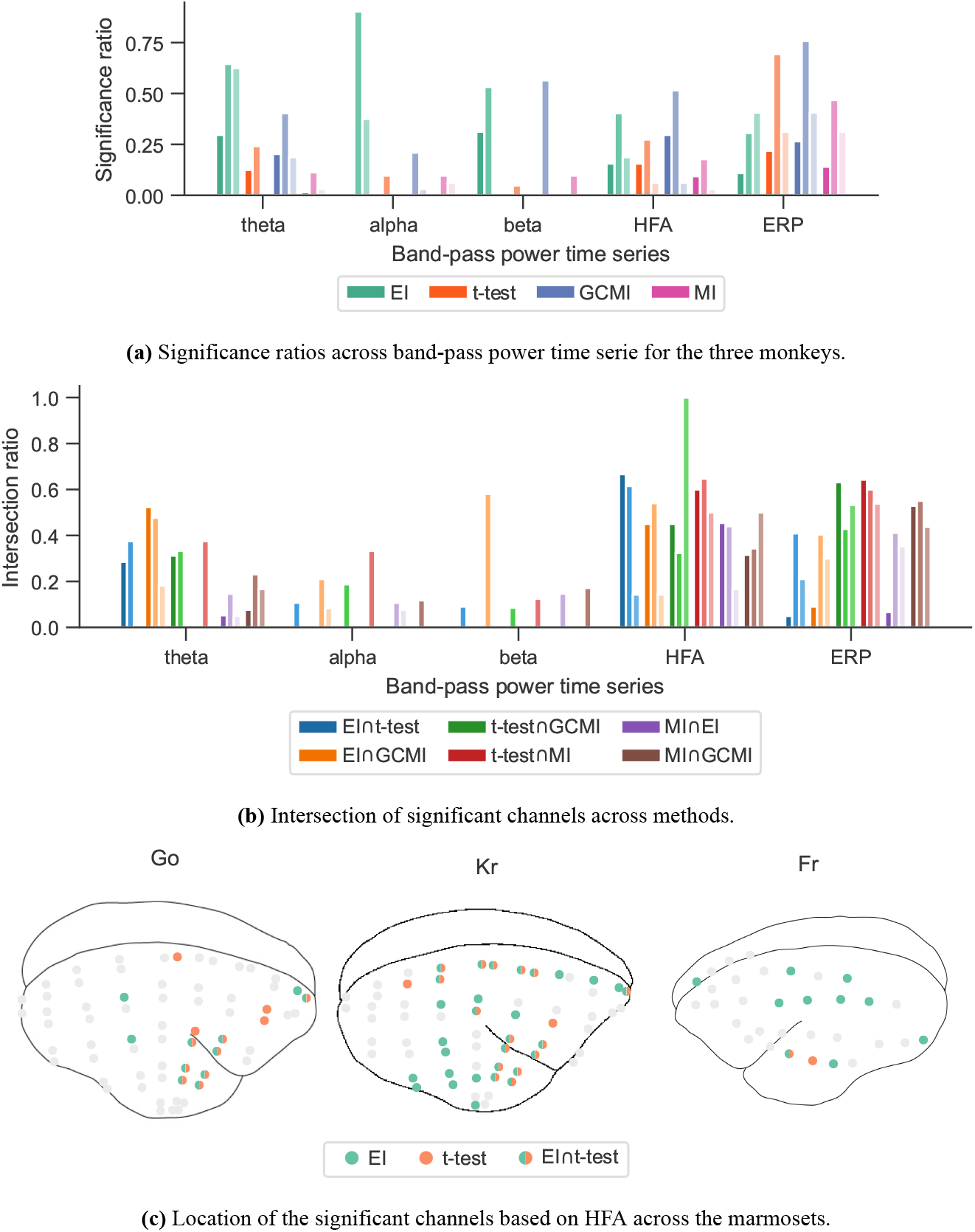
Performance of the different methods for the Roving Oddball paradigm across the three marmoset monkeys “Go”, “Kr” and “Fr”. **5a**: Global significance ratio for each marmoset. **5b**: Intersection of the significant channels for each method combination. Each number is shown relative to the total number of significant channels for each method combination. **5c**: Location of the significant channels for EI and t-test for HFA (monkey “Go” has 64, “Kr” 62, and “Fr” exhibits 32 channels).

**Figure 6:**
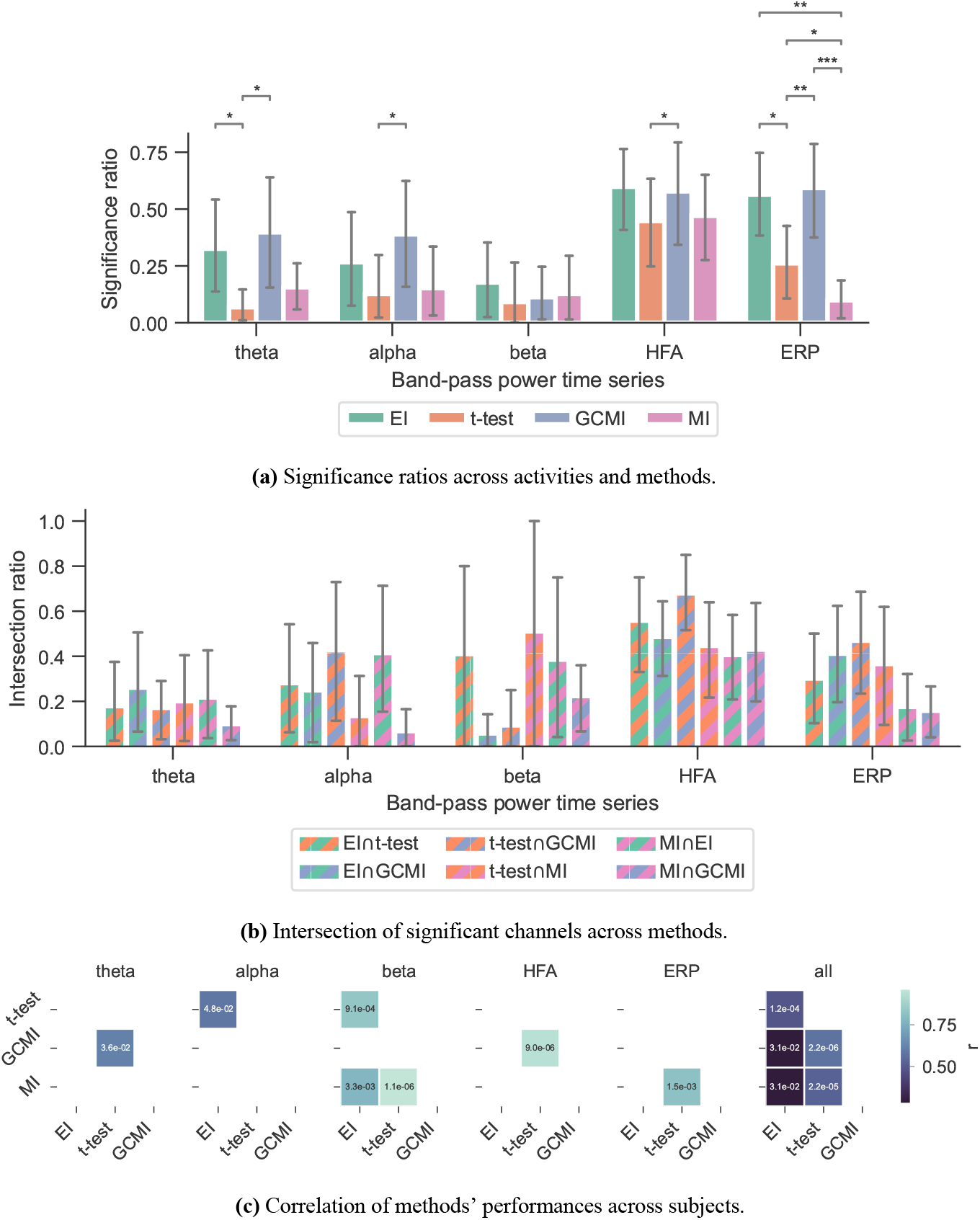
Performance of the different methods across subjects for the vWM task. **6a**: Performance of the different methods. The error bars indicate the 95% CIs across subjects. **6b**: Significance ratio across representation. Statistical significance is indicated with * p≤5*e*−2, ** p≤1*e*−2, *** p≤1*e*−3 and **** p≤1*e*−4. **6c**: Correlation matrices comparing the subject-specific significant ratios. The respective p-value is annotated in each square.

**Figure 7:**
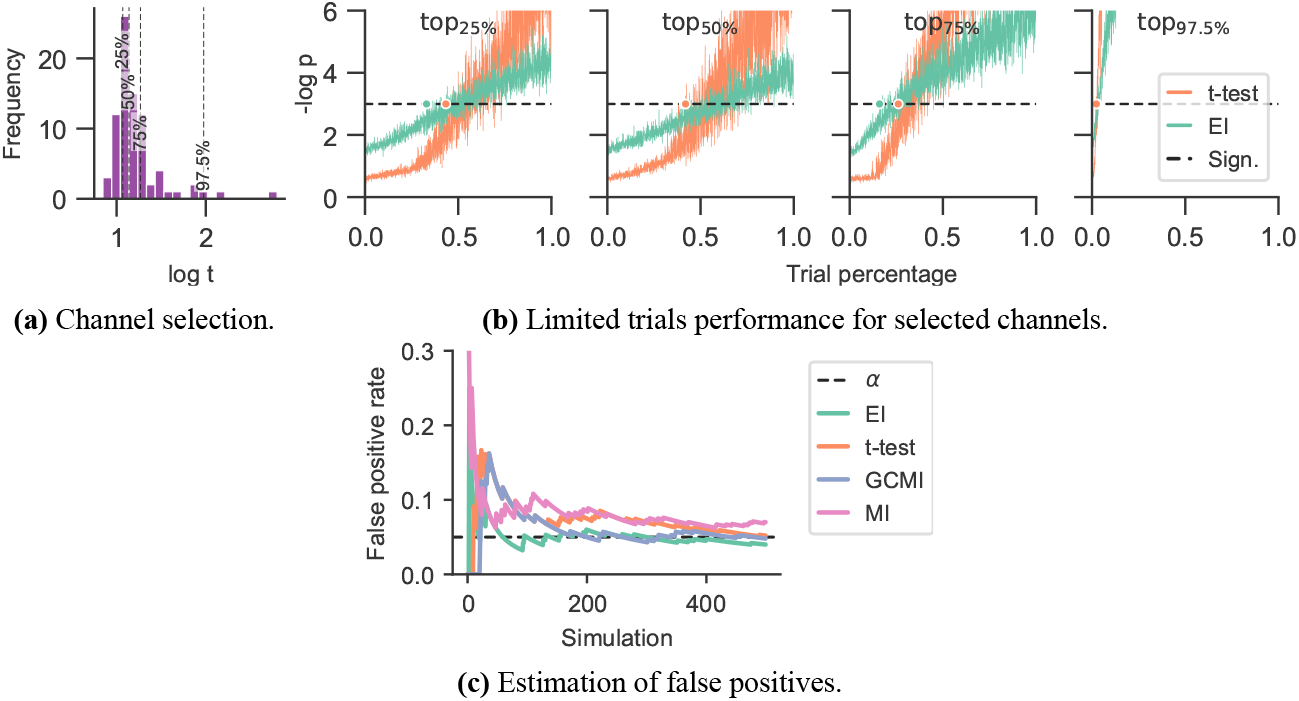
Performance in the case of a limited amount of data and estimation of false positives. **7a**: Selection of channels according to their respective t-value. The channels emerged from HFA or the Optimum-1 paradigm and were located in the respective 25, 50, 75, and 97.5-percentiles of the t-value distribution. **7b**: The number of trials was reduced by randomly selecting a varying number of trials for each condition. This was followed by repeatedly applying the measures (50 times for each percentage). The dashed line represents the significance threshold (p=0.05). Note that the y-range is limited to 6 (corresponding to a p-value of 2.5*e*−3). 100 % of the trials was around 759.50 ± 360.85 trials for deviant responses and 715.14 ± 388.12 trials for standards for each channel. The dots on the significance line indicate the trial percentage when each method exceeded the significance level. **7c**: False positive rate estimation by repeatably discriminating two random samples with 100 observations each, drawn from the standard normal distribution. The significance level *α*=0.05 is indicated with a black dashed line.

### Optimum-1 Paradigm

Across the 1078 channels stemming from the Optimum-1 paradigm, for alpha and HFA signals, EI’s significance ratio was significantly greater than MI or t-test (Fig. 3a) while the t-test showed the greatest significance ratio for the ERP (two-sided paired t-tests on a subject level, p=[1.02e−2, 4.17e−2, and 3.666e−3]). MI resulted in the lowest ratios, although it was only significantly lower on a subject level for HFA (two-sided paired t-test, p_max_=4.17*e*−2). Note that for human iEEG, the number of channels and their distribution is not constant across subjects and can thus vary greatly. Consequently, the fewer channels for a subject, the less robust the estimated individual significance ratio.

Given the differing sensitivity to deviating sounds across individual brain regions and the number of participants for the Optimum-1 paradigm, we then compared the methods’ significance ratios across individual cortical areas. For this reason, we evaluated the methods’ performances across different areas comprising temporal, frontal, insular, peri-central sulci, and anterior cingulate cortices (ACC), as well as the hippocampus (Fig. 4; see 5, for exact definitions). Overall, the significance ratios of EI aligned with the other methods across brain regions and representation. Specifically, the ratio for higher cortical areas such as the superior or middle frontal cortex was low while being higher in areas such as the superior temporal plane (which includes Heschl’s gyrus). The NFT method together with MI showed the lowest sensitivities across brain regions. However, there were higher or equally high ratios in responsive brain areas such as the superior temporal plane or posterior insular cortices.

### Performance of NFT

For the Optimum-1 paradigm, we further evaluated the NFT of all contact sites. The number of responsive channels, i.e., the number of channels that solely tagged the frequency of standards or deviants, was lower in comparison to all other methods (two-sided paired t-test between NFT and MI, p=3.55e−2).

### Roving Oddball Paradigm

The observations from this data set are in line with the results from the marmoset recordings, which had 160 channels in total. Based on the significance ratio of each marmoset monkey, for theta, alpha, and beta signals, EI detected more channels than the other approaches. For HFA, GCMI and EI performed similarly well, while for ERP, GCMI and t-test performed best (Fig. 5a). Additionally, common to all methods was the variability in significance ratios across monkeys.

### vWM Task

A similar observation was made by considering EI’s performance for the vWM task (Fig. 6a). Across all band-pass power time series, EI showed the highest significance ratios. Interestingly, for the ERPs, both EI and GCMI had a significantly greater significance ratio than the t-test (p_max_=8.21*e*−3). Besides that, especially GCMI exhibited a consistently higher significance ratio across band-pass power time series in comparison to the Optimum-1 and Roving Oddball paradigms.

### Method Overlap Across Data Sets

To assess channels that the methods commonly detected, we divided the number of common significant channels (i.e., the intersection) by the total amount of significant channels for each pairwise method combination (Fig. 5c, i.e., each method’s specific unique set of significant channels plus their intersection) and defined this ratio as the intersection ratio. For the Optimum-1 paradigm, the intersection ratio between the t-test and GCMI was the greatest across all band-pass power time series (Fig. 3b). For beta and HFA signals, there was also a high intersection ratio between the t-test and GCMI. However, for these band-pass power time series, the number of significant channels was low (Fig. 3a), which led to higher percentages. This can also be observed by correlating the method-specific channel distributions across subjects (Fig. 3c), i.e., by correlating the significance ratios across subjects for each method-to-method combination. Besides that, the approaches correlated the least for beta signals, which were also the signals with the lowest significance ratios. Overall, there was a high correlation across methods (Fig. 3c).

The relatively high intersection ratio between the t-test and GCMI across band-pass power time series was not observable for the Roving Oddball paradigm and the vWM task. For the former, the intersection ratios were the highest for HFA and ERP signals (Fig. 5b), while for the latter it was highest for HFA (Fig. 6b&6c). Furthermore, the same effect as for the Optimum-1 paradigm recordings occurred for the Roving Oddball paradigm. For HFA, the monkey Fr showed maximum intersection ratios between t-test, GCMI, or MI. Of note, the respective significance ratios were rather low (Fig. 5a). Moreover, when comparing Kr to Go it appeared that the latter had a smaller number of significant and overlapping channels for EI and t-test. However, because Kr exhibited more significant channels for both methods, the intersection ratio was of comparable magnitude for both monkeys.

### Reduced Trials and False Positives

Considering the two best-performing methods of EI and t-test, when reducing the number of trials they only marginally differed in reaching statistical significance although EI tended to reach this threshold slightly earlier (Fig. 7b). In terms of the simulation-based estimation of the false positive rate estimates, all methods showed similar rates which were close to the level of significance *α* of 0.05 (Fig. 7c).

## Discussion

Despite its ability to estimate the absolute information contained in individual brain responses, the use of AIT in neuroscience is limited. Hence, information-theoretical approaches in neuroscience are needed to facilitate its use. With the measure of EI, we proposed information-based task condition discrimination as an alternative to the classical t-test and other information theory approaches. Using compression as the core principle, this procedure quantifies the similarity between recordings stemming from different conditions. We applied all procedures to event-related potentials extracted from different frequency bands in three iEEG datasets and compared it to that of t-test, MI, GCMI, and NFT. We discuss the performance and sensitivity of these methods, their pros and cons, and similarities and differences.

### Sample versus Mean Response Approach

Besides their individual procedures in estimating test statistics, the examined methods also differ in how the data were processed. While for the t-test and GCMI approach, each sample or time point across trials is considered independently, EI and MI in its current implementation, make use of all time samples by computing the respective mean responses across trials. Consequently, for each channel, the first two methods result in test statistics along the time axis, while the last two output a single value channel-dependent test statistic. Such a time course is advantageous when the core interest is within the time domain or latency of responses, i.e., at which time point responses differ most. However, when it comes to assessing how sensitive contact sites are towards different conditions, this time course is of a minor role. To assess a channel’s significance, it is necessary to reduce the time course of the test statistics to one single value. Additionally, the complete time series of the respective statistic also needs to be corrected for multiple comparisons across channels and trials (i.e., correction of a 2D array that covers samples and channels versus a 1D array that only includes channels). Therefore, corrections might undermine the sensitivity of sample-based approaches.

### Performance of EI

Adopting the sample-based approach mentioned above is not beneficial for EI. While MI or GCMI estimates the underlying distribution of the input data, EI is a compression-based approach. It thus compresses the entire time series while exploiting structures along the time axis. This mechanism is a key feature of this approach and is possibly responsible for the higher significance ratios across evoked potentials and other band-pass power time series. If EI is applied along trials for each sample, this feature of exploiting structures along the time axis would be removed because the input would be a concatenation of different signals. Further, especially for time course responses that are close in magnitude or partially overlapping, i.e., situations where the classical t-test might fail to disentangle condition differences measuring EI proves beneficial. By exploiting complementary structures within both responses, subtle information-grounded differences that extend beyond the width of one sample can be identified in such scenarios. Additionally, the binning parameter or resolution along the time axis plays minor importance when it comes to computation time, while it can have a great effect on the computation time of GCMI or cluster-based t-tests.

Especially for the beta and HFA power time series, EI appears to be a useful tool for detecting active channels (Fig. 3a). Across all data sets, EI exhibited the highest significance ratios compared to the t-test, MI, and GCMI. HFA presumably carries stimulus mismatch or prediction error signals (26). In that regard, detecting a high number of channels in HFA discriminating between standard and deviating sounds in regions such as the hippocampus appears to be especially interesting (Fig. 4 and 6a).

### Performance of t-test

For the ERP, the t-test showed the greatest significance ratios for the Optimum-1 and Roving Oddball paradigm. One possible reason for this could be the generally low variance (or standard error) in EPRs compared to the other electrophysiological representations. This low variance leads to statistically strong differences in ERP amplitudes resulting from the different stimulus conditions. Further, the number of samples that significantly differ in amplitude across conditions after correcting for multiple comparisons can be rather low for a channel (i.e., the entire trial versus only a few sample points differ). EI, on the other hand, considers the full-time course and only assesses channels as significant when the information content of the entire mean responses differs. However, the high performance of the t-test is not observable for the vWM task. Here, both EI and GCMI detected a greater number of responsive channels, while also showing a relatively high intersection ratio between t-tests, respectively.

### Performance of NFT and MI

Compared to all the other measures (EI, t-test, MI, and GCMI), the NFT approach had inferior performance. However, when considering individual ROIs, it performed equally well for regions close to or belonging to the temporal cortex. Notably, NFT is based on a different procedure. It exploits the static presentation rates of stimuli visible in the spectral decomposition of the entire sequence. Fig. 2 shows such a “tagging” effect in an exemplary channel, where a specific brain region synchronizes to the exact frequency of the stimuli presentation. The highest SNR or power was contained at 1.92 Hz, which was the presentation rate of the tones (standard and deviants). In addition, it also showed a peak at half this frequency, which is the combination of either standard-to-standard or deviant-to-deviant tones. This implies that this brain region discriminates between standard and deviant tones. Importantly, one pre-requisite for the frequency tagging approach is that the presentation rate is static. Any temporal jitter during the presentation critically disturbs the “tagging” effect. A possible reason for the relatively low performance could be related to this unique feature of NFT combined with the employed multi-featured oddball paradigm. It may perform better in paradigms with fewer deviant types than in the present task containing eight types of deviant tones.

When it comes to performance, MI was close to NFT. It detected fewer channels than the other methods, especially for alpha, beta, and HFA band-pass power time series. It is important to notice that for comparison reasons, we implemented MI in the same way as EI. That is, the entire time series of a mean response was considered. It is also possible to implement it in the sample-based fashion of GCMI, where it has been shown to operate similarly well (3).

### Method Overlap

Overall, the sample-based approaches of t-test and GCMI correlated the most (Fig. 3c). Besides that, there was no clear outlier method apparent for the analyzed data sets. The same conclusion can be drawn considering the significance ratios across individual brain regions (Fig. 4). This can be seen as a validation of the EI measure: While showing higher significance ratios in some ROIs (e.g., hippocampus or precentral sulcus), it performed similarly well as the other methods across brain areas and did not indicate implausible results. Further, our simulation-based analysis indicates that none of the methods exhibited a higher number of false positives than the others (Fig. 7c). All methods’ false positive rate estimates converged to the theoretical value of 0.05 pointing towards the conclusion that all methods are equally reliable.

### Choice of Number of Bins

Furthermore, the number of bins along the potential or power dimension during the binning procedure was held constant across data sets. We chose 128 bins for EI and four bins for MI (3, 5, 10). Adapting this parameter might be needed when the relevant trials are of increasing length, i.e., it might be necessary to reduce the number of bins for the EI measure for longer-lasting trials. The main motivation for this step is to simplify the signal to an extent that ensures that the compressor keeps operating effectively. However, both EI and MI should be robust to this parameter choice. For example, EI showed similar results when varying the number of bins from 128 to 64, but showed a decreased performance when using only 32 bins (6, 7, 10). An alternative to adapting the number of bins is to segment the time series into shorter segments, which would then yield segments of EI values along the trials, or to down-sample the signal along the time axis.

### Reduced Number of Trials

Lastly, in the case of limited data availability, the two best-performing methods showed relatively similar performance (Fig. 7). For lower t-values (below the 75-percentile), both methods needed around 50 % of the maximally available trials to identify a channel as event-responsive. However, EI appeared to require slightly fewer trials than the t-test. Noteworthy is that for channels with relatively distinct responses to standards and deviants (above the 97.5-percentile), both measures only required around 3 % (ca. 22 trials) of the available trials to identify the respective channel as significant.

### Performance Across Paradigms

Altogether, the examined neurophysiological data sets stem from two species (human and non-human primates) and employ both passive and active paradigms as well as auditory or visual stimuli. In contrast to the Optimum-1 and Roving Oddball paradigms, in the vWM study subjects were instructed to solve a memory task where they were exposed to visual stimuli. Moreover, as opposed to comparing standard with deviant tones responses, for the vWM paradigm, the probe period was compared to the baseline period based on one cortical region. Regardless of these differences, EI as well as MI, and GCMI performed robustly across all data sets (Fig. 3a, 5a, and 6a).

## Conclusion

Taken together, our findings demonstrate that EI, MI, as well as GCMI, constitute viable approaches to discriminate differences in neurophysiological recordings of evoked responses. Specifically, EI competed well in detecting iEEG channels sensitive to deviating sound types across diverse types of electrophysiological responses among the methods considered. Especially for beta and HFA signals, EI detected a higher number of sensitive channels in comparison to the other procedures. Future studies focusing on EI could explore this further by extending it through a hybrid approach, i.e., using Shannon’s information theory besides AIT. Another possibility is to focus on the compression method. Modern compressors such as LZMA, Brotli, or neural network compressors might be able to increase EI’s performance. On a more general note, information-based encoding proved to be a worthwhile tool in assessing where in the brain neural responses differ across experimental conditions.

## Author Contributions

JF, KG, AKS, TE, and AB contributed to the conception and design of the study. AB and AL carried out the experiment and collected the data. JF, AB, KG, TE, and AKS contributed to the interpretation of the results. JF wrote the manuscript with inputs from AB, KG, TE, and AKS. All authors revised the manuscript. All authors read and approved the final manuscript.

## Ethics Approval and Consent to Participate

This study was approved by the Research Ethics Committee of El Cruce Hospital, Argentina, the Regional Committees for Medical and Health Research Ethics, Region North Norway, and the University of California, Berkeley. Patients gave written informed consent before participation. Electrode placement for the human iEEG recordings was solely determined based on clinical considerations.

## Acknowledgments

We thank the patients for kindly participating in the studies. We want to express our gratitude to the EEG technicians at El Cruce Hospital and Oslo University Hospital-Rikshospitalet for their support. We thank Pål G. Larsson, Jugoslav Ivanovic, and Ludovic Bellier for their collaboration. Further, we would like to thank Misako Komatsu, Kana Takaura, and Naotaka Fujii for publicly sharing their data.

## Data Availability Statement

Custom analysis codes written in Matlab are available at osf.io/tnvc4. Neurophysiological data for the roving oddball paradigm is publicly available at neurotycho.org/auditory-oddball-task. Due to the confidential nature of the data, the patients’ datasets analyzed for the Optimum-1 paradigm study are not publicly available. The ethical approval conditions do not permit public archiving of study data. Readers seeking access to the data should contact the AB, Department of Psychology, University of Oslo; the Research Ethics Committee of El Cruce Hospital, Argentina; and the Regional Committees for Medical and Health Research Ethics, Region North Norway. Requests must meet the following specific conditions to obtain the data: a collaboration agreement, a data-sharing agreement, and formal ethical approval. Data for the vWM task can be made available from AL upon reasonable request.

